# Sequence evidence for common ancestry of eukaryotic endomembrane coatomers

**DOI:** 10.1101/020990

**Authors:** Vasilis J. Promponas, Katerina R. Katsani, Benjamin J. Blencowe, Christos A. Ouzounis

## Abstract

Eukaryotic cells are defined by compartments through which the trafficking of macromolecules is mediated by large complexes, such as the nuclear pore, transport vesicles and intraflagellar transport. The assembly and maintenance of these complexes is facilitated by endomembrane coatomers, long suspected to be divergently related on the basis of structural and more recently phylogenomic analysis. By performing supervised walks in sequence space across coatomer superfamilies, we uncover subtle sequence patterns that have remained elusive to date, ultimately unifying eukaryotic coatomers by divergent evolution. The conserved residues shared by 3,502 endomembrane coatomer components are mapped onto the solenoid superhelix of nucleoporin and COPII protein structures, thus determining the invariant elements of coatomer architecture. This ancient structural motif can be considered as a universal signature connecting eukaryotic coatomers involved in multiple cellular processes across cell physiology and human disease.

## Main Text

Nuclear pore complexes (NPCs) are modular assemblies embedded at the points of fusion between the inner and outer membrane of the eukaryotic nucleus that mediate nucleocytoplasmic transport ^1^. The overall architecture and composition of the NPCs is largely taxonomically conserved, indicating early origins in the eukaryotic tree ^2^. In particular, nucleoporins including those at the outer ring coat forming the Y-complex (outer ring coat Nups or Y-Nups) share certain key structural and architectural similarities, possibly due to deep divergence ^3,4^. These features extend beyond the nuclear pore, namely the COPII coat associated with anterograde transport from the rough endoplasmic reticulum to the Golgi apparatus and the COPI coat associated with the reverse, retrograde transport ^5^, suggesting a common origin of endomembrane coatomers, one class of which is represented by the NPC coat ^6^.

The divergence of nucleoporin families has been proposed on the basis of global structural but no specific sequence evidence, especially for the Y-Nups ^7^. This presumption is based on detailed structural analysis, presence of beta-propeller repeats at the N-terminus, an alpha-solenoid superhelix at the C-terminus and other architectural elements with regard to the multi-domain composition of Y-Nups ^8^. In particular, the solenoid superhelix of the resolved structures for Nup75, Nup96, Nup107 (Y-Nups) and Nic96 – reminiscent of the tetratrico-peptide repeat (TPR) domain ^9^ – is also present in Sec31 and Sec16, building blocks of the COPII vesicle coat ^10,11^. Much attention has been paid to this structural element as a common architectural motif across diverse coatomer molecules, and has thus been named Ancestral Coatomer Element 1 (ACE1) ^10,12^, favouring the hypothesis of deep divergence over convergent evolution ^13^. Yet, no sequence signature for ACE1 has ever been detected, either for Y-Nups/Nic96 or Sec31/Sec16, while the structure determination and comparison of these coatomers revealed this surprising structural similarity ^10^. ACE1 might be considered as a structural manifestation of the likely common origin of NPC and COPII coats ^12^, but has never been observed outside these complexes ^14^. A combination of phylogenomic profiling and structural predictions has further extended this relationship to the intraflagellar transport complex (IFT) of the cilium ^15^, across eukaryotic phyla and their representative genome sequences ^16^. Affirming an earlier hypothesis for the homology of the IFT complex with endomembrane coatomers ^17^, IFT-A components IFT122, IFT144/WDR19 and WDR35 and IFT-B components IFT172 and IFT80 are detected as ancestrally related to COPI subunits, yet without a connection to nucleoporins or a reference to the ACE1 structural motif ^16^. Therefore, despite abundant sequence and structural data for this motif, the identification of ACE1-containing molecules and their relatives remains highly challenging, a task partly achieved only by a mixture of sequence profiles, alpha helical predictions and domain architecture considerations ^10^. Herein, we present sequence evidence for the long suspected common origin of NPCs, COPIIs, IFTs and other coatomer systems across eukaryotes, unifying previous insightful hypotheses and detailed structural studies ^18^.

In our quest for multi-domain architectures for the structural and functional analysis of the Y-complex ^19^, we have encountered a unique, subtle sequence similarity with a critical, missing link between the Nup75 sequence profile and the Nup98-96 (Nup96) sequence of the insect species *Harpegnathos saltator* (GI:307191801) ^20^. Thanks to the recent availability of genomic information across many eukaryotic genomes, gaps in this particular region of genome sequence space are being filled rapidly by homologs which can connect hitherto seemingly unrelated protein sequence families – in this case Y-Nups, via significant sequence similarities (see also Supplement). We have further pursued a rigorous analysis of this puzzling connection between Nup75 and Nup96 families by conditional iterative sequence profile searches, using the Nup75 sequence profile as a query – Nup75 alignment positions 339-2024 in DS03 of our previous report ^19^ (Data Files S1-S4). By inspecting thousands of alignments, we were able to detect sequence signals of the divergent alpha-solenoid superhelix within 3,502 sequences in the non-redundant protein database (effective date December 2013) (Figure S1). Having initially excluded Nup107 (which terminates the search early, see Methods in Supplement), we uncover the deeply divergent sequence relationships between Nup96, Sec31, WDR17, Nic96, IFT140 (from IFT-A), IFT172 (from IFT-B) and finally Nup107 in this order (Figure 1) – while IFT144/WDR19 and IFT122 (IFT-A components), Sec16 and Clathrin (but not COPI) are marginally detected beyond the set threshold, thus unifying NPC, IFT, COPII, and Clathrin ^21,22^, as well as uncharacterized molecules such as WDR17 (see Supplement).

**Figure 1:**
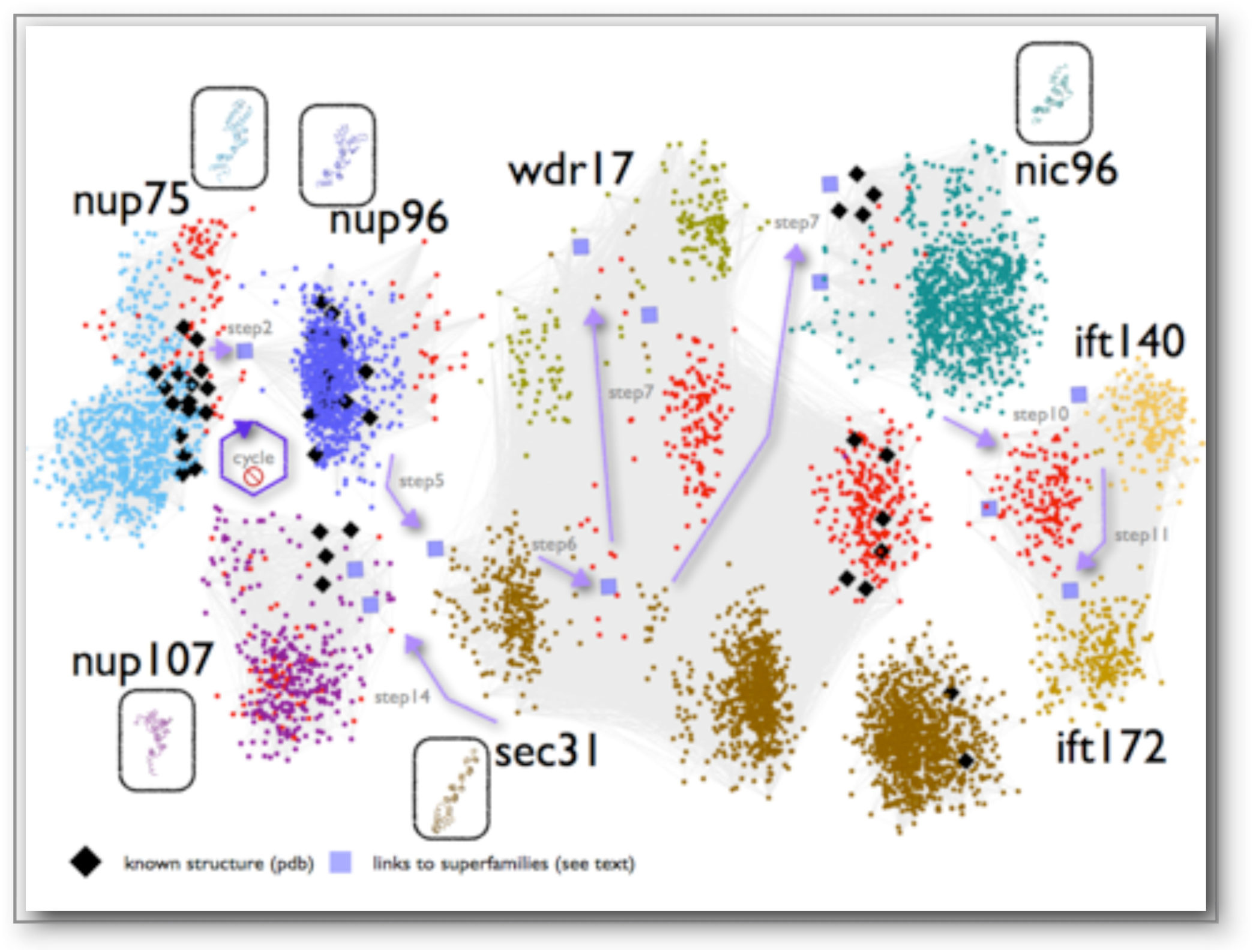
Pictorial representation of the sequence space walk connecting components of the nuclear pore complex, COPII and intraflagellar transport. Each family is represented by a distinct color; exceptions are homologs of known structure depicted as black diamonds and previously uncharacterized (unannotated) protein sequences depicted as red dots. Proteins of known structure representing specific families are depicted in oval boxes with identical colors, with identical orientations (as in Figure 2). Inter-family connections detectable by sequence searches are represented by thin light grey lines (see Methods). Intra-family connections revealed by iterative profile searches – otherwise undetectable, are depicted by light purple arrows for the corresponding steps and squares for sequence links across superfamilies. The cycle with a ‘stop ’ sign refers to the exclusion of Nup107 family which terminates the search early. Only a representative subset of hits is shown for clarity. An annotated version of this two-dimensional layout is available as Data File S11.

**Figure 2:**
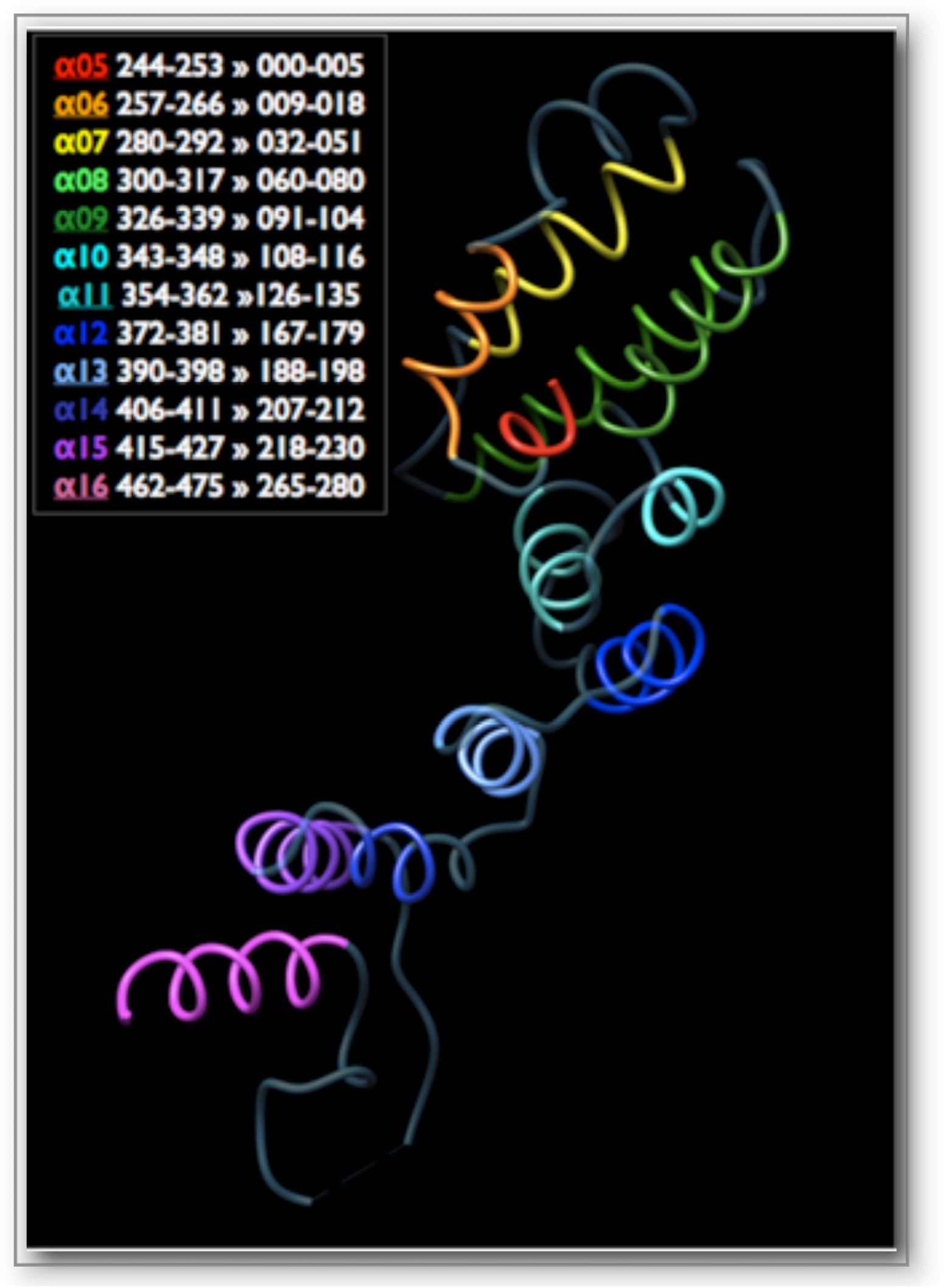
Sequence conservation across endomembrane coatomer structure components. Figure 2a: The Nup75 structure (3F3F_C) corresponding to the detected ACE1-like alpha-solenoid superhelix motif is shown. Individual helices α5-α16 are colored by unique colors, warm colors representing the most conserved segments (α5-α6, α15-α16). Sequence positions for each helix are shown in the legend, according to the comparative structural analysis of ACE1 ^10^, followed by the corresponding positions in the alignment in Figure 2b. This orientation is used throughout this work, corresponding to the crown and the second half of the ACE1 trunk – see also *Figure 5* in ^10^.

**Figure 2b:**
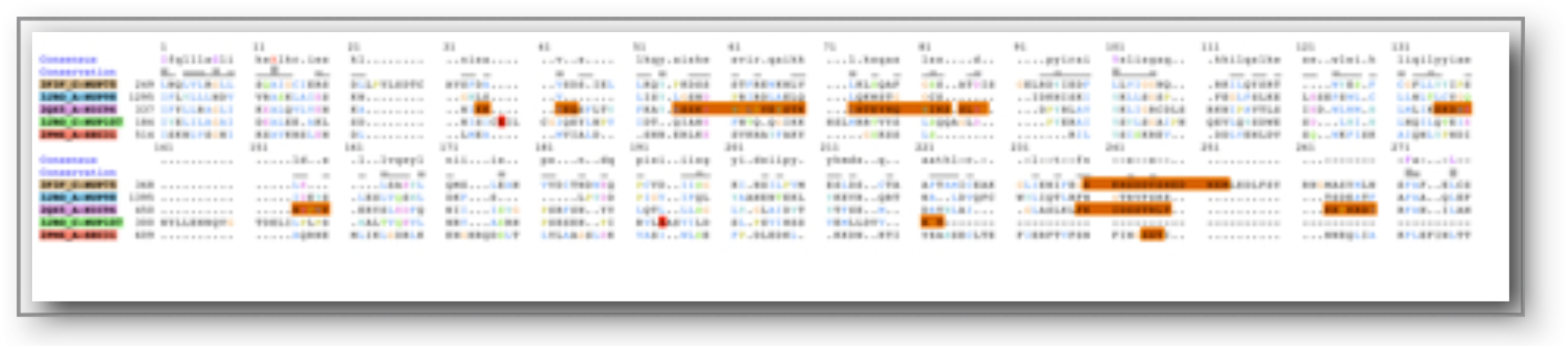
Sequence alignment of five alpha-solenoid superhelix motif-containing representative structures. Aligned positions are established by direct comparison of the KMAP-13 profile against the structure database. PDB codes are given, followed by the description of the corresponding protein chains. The total length of the alignment is 280 residues. A consensus sequence and a skyline conservation plot are provided. Regions missing in the structure database entries are shown in orange. See Data File S9.

**Figure 2c:**
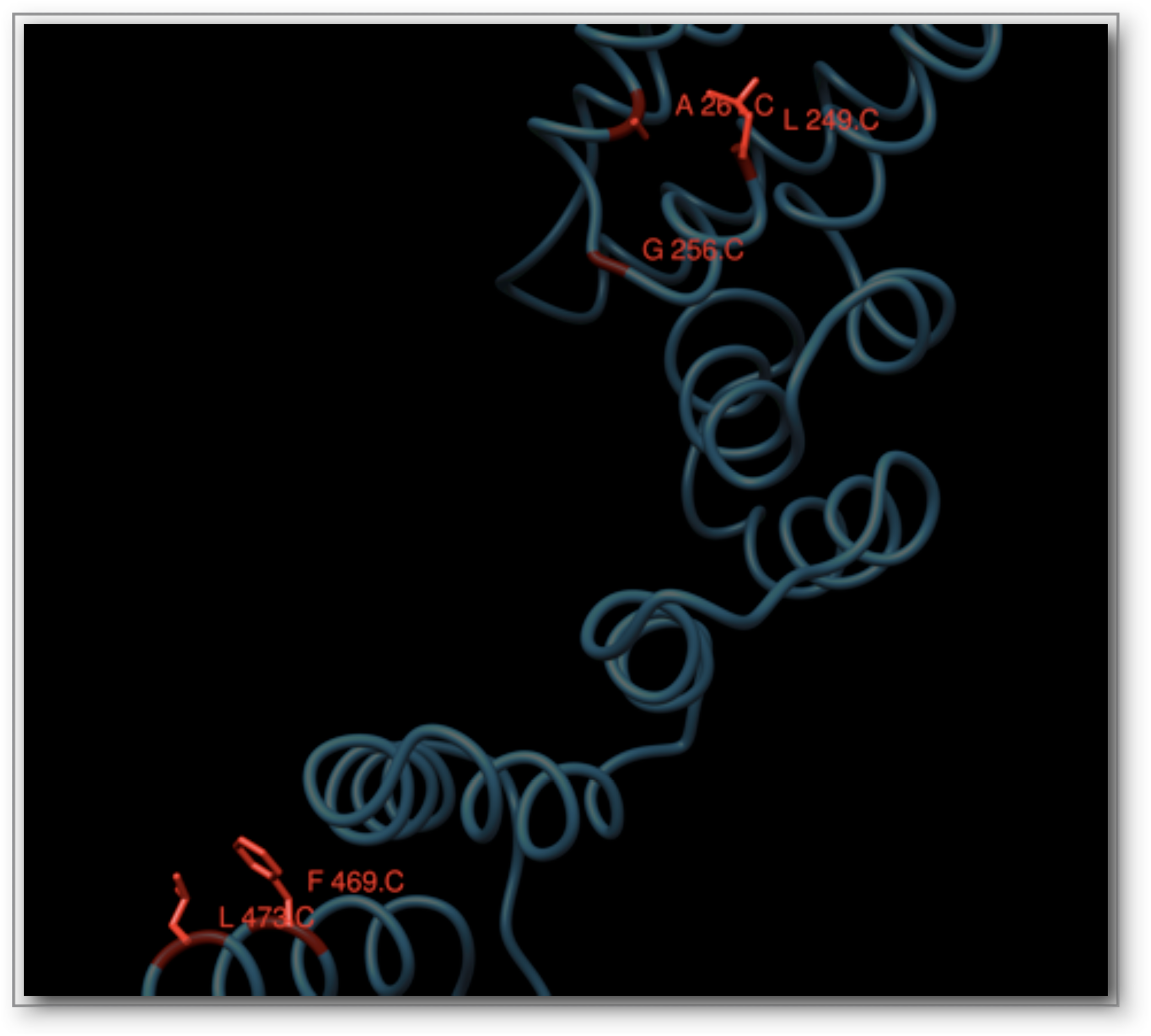
Structural context of the evolutionarily conserved positions in Nup75. N-terminal Leu-249, Gly-256 and Ala-261 (C for chain C of 3F3F) (see Figure 2a) and C-terminal Phe-469 and Leu-473 correspond to alignment positions I1, G8, A13 and F272, L278 respectively (Figure 2b). The C-alpha trace is shown in dark blue and the subset of conserved positions is shown in red, along with the side chain representations. Alignment position L101 is not highlighted as it is frequently substituted by a tyrosine residue (Figure 2b).

This complex sequence profile search has been crucially based on the manual exclusion of 15 putative false positive cases (and, hence, of their homologs in future searches) (Table S1), some of which tend to appear in our profile sequence queries more than once. At each step, this procedure – which can be regarded as a genuine *sequence space walk* – unravels specific subsets of increasingly distant homologs in a highly controlled manner (Figure S1, Table S2). To ensure reproducibility, we have carefully repeated and documented these profile searches, until the process encounters noise, i.e. spurious sequence similarities for which no evidence of ACE1-containing motifs is available either in annotation records or reverse sequence searches (Data Files S5). A visual representation of an increasingly sensitive sequence profile across sequence space is provided as a video file (Video S1). To assess coverage, we have also interrogated the protein database using Entrez^®^ text queries, and retrieved 5662 redundant entries (61 duplicate, 5601 unique), many of which, however, represent false positive identifications of the corresponding motifs (by automatic assignment) (Data File S6). To maintain precision at virtually 100% (Figure S1), as indicated by detailed structural validation and interpretation (see below), coverage is somewhat compromised at this particular database search significance threshold and is indeed under- estimated, while in principle could be increased by imposing length constraints similar to sequence searches. The iteratively derived profile named KMAP-13 for ‘euKaryotic endoMembrane ACE1 Profile at Step 13’ is made available (Data File S7), along with the hit table containing sequence identifiers, to facilitate the extraction of the corresponding database entries and future updates (Data File S8).

A key result of this sequence space exploration is the demonstration that three nucleoporin families – namely Nup75, Nup96, Nup107 – not only share structural similarities but these similarities arise from divergent evolution at very low sequence identity levels (minimum sequence identity across runs 3-9%, average 6.8%) with statistically-significant alignments (p<0.001). Our sequence profile searches unambiguously underline the deep phylogenetic connection of these Y-Nups, as well as Nic96 and Sec31 (see Supplement). In this well-defined, newly discovered sequence space locality of these homologous molecules containing the alpha-solenoid superhelix, there are five protein families represented by resolved three-dimensional structure homologs, namely Nup75, Nup96, Nup107, Nic96 and Sec31 (Figure 1) – the corresponding Sec16 region is also correctly detected, albeit below threshold. To validate the sequence profile-driven alignments, we superimposed the four known nucleoporin as well as Sec31-COPII structures on the basis of aligned positions for five conserved residues: the structural superposition verifies our results, as four structures are superimposed precisely along the alpha-solenoid superhelix with RMSD values <3Å (Figure S2, except Sec31’s last helix hairpin) – with better fit towards the N-terminal part of the ACE1 alpha-solenoid. Surprisingly, this is the first time that sequence information alone strongly reflects the structural similarity of these molecules as previously observed ^12^, thus both delineating the evolutionary history of ACE1-like motifs and supporting the hypothesis that coatomer systems arose by divergent evolution ^23^ (Figure 2). Furthermore, the structural partitioning of ACE1 and relatives into crown (α5-11), trunk (α1-3,α13-19) and tail (α21-28) can now be viewed from an evolutionary perspective, where helices α5-16 across the crown and trunk segments represent the conserved core of ACE1 (Figure 2a). In a length of 280 residues, there are only three invariant positions linked by divergence: Ala (A13), Phe (F272), Leu (L278) (Figure 2b); a number of other conserved positions are also observed namely Ile (I1), Gly (G8), Tyr (Y101) – both these sets are used for structural superposition. It should be noted that the full alignment unravels limited variation across these positions, for example A13 is 84% present sporadically substituted by Ser or Thr and F272 is frequently substituted by Tyr (Data Files S5).

The deep connections brought to light by this sequence space walk resolve a long-standing issue of coatomer phylogeny across eukaryotes and permit the evolutionary dissection of rich structural data for ACE1 alpha-solenoids (Figure 2). Remarkably, helix α8 of Nup75 ^12^, corresponding to helix α7 of ACE1 ^10^, does not exhibit conservation across those families as previously reported ^12^. The most conserved block of ACE1 resides towards the N-terminal part of the crown, namely helices α5-α6 (positions 1-22, Figure 2b-c). The alignment quality decreases towards the C-terminal part, with the exception of invariant positions 272 and 278 (undetectable in the Nup107 structure – 3jroC). The conserved ACE1 block ranges between invariant alignment positions I1 (α5 position 11; Ile in other structures, Leu-249 in Nup75) and A13 (Ala-261) maintaining the hairpin α5-α6 contact (Figure 2c), while position G8 (Gly-256, Asn in Nup96) acts as helix breaker for α5. At the C-terminal region of the conserved ACE1 block, position F272 (Phe-469) packs against position L278 (Leu-473), stabilizing α16 with α15 of the trunk, and possibly α1 as well (Figure 2c). We reason that these non-polar residues might not contribute towards interface contacts and are most likely involved in maintaining the ancestral structural integrity of ACE1, despite astonishing variation acquired elsewhere in this structural motif ^24^. This is a testable prediction that could be validated by assessing the impact of evolutionarily conserved regions ^19^ for the stability of endomembrane coatomer components represented by the currently available structures as above.

The evolutionary dissection of endomembrane coatomers exhibits a strong structural conservation of the alpha-solenoid superhelix with family-specific sequence variation and adaptive association with beta-propeller motifs in the case of the Y-complex, e.g. Nup75 with Seh1 and Nup96 with Sec13 ^8^. A particular instance of the coatomer superhelix shared between nucleoporins and COPII components has been attentively termed ACE1 based on structural similarities ^12^. Our analysis provides specific sequence evidence for the deep divergence of three Y-Nups/Nic96 and COPII, as previously proposed on structural grounds ^18^. We further extend the presence of the coatomer superhelix to some of the longest components of the IFT, namely IFT140 and IFT172, recently predicted as remote relatives by phylogenomic analysis ^16^. The detection of a number of IFT core components ^25^ by this sequence space walk suggests that they play a key role in the ciliary pore complex (CPC) that regulates transport ^26^, analogously to the NPC ^27^. Further structural analysis of IFT components, so far achieved for a number of smaller IFT core proteins ^28^, can unravel the alpha-solenoid superhelix in some of the longest IFT members, such as IFT144 or IFT172, involved in human skeletal ciliopathies ^29^. The identification of WDR17 points to its involvement in eye gene expression and possibly disease ^30^, further supported by positive selection pressure in dolphin sensory systems ^31^ and by proteomics detection as a conserved element in Joubert Syndrome-associated ciliary signaling subdomain ARL-13 ^32^. The compelling sequence similarity across coat complexes and associated processes enhances proposals about a common origin of endomembrane coatomers early in eukaryotic evolution ^33,34^. The puzzling connections revealed by deep divergence of coatomers offer new perspectives for their emerging implication in coupling multiple cellular roles, such as the kinetochore involving the Y-complex ^35^, nucleoporins associating with histone-modifying complexes ^36^ and the centrosome connecting to the nucleus, the Golgi apparatus and the eukaryotic cilium ^37^.

Supplement

37+26 additional references

## Acknowledgments

We thank Manuel Irimia (Centre for Genomic Regulation, Barcelona, Spain), Daniel Lundin (SciLifeLab, Stockholm, Sweden) and Kai Simons (Max Planck Institute for Molecular Cell Biology & Genetics, Dresden, Germany) for comments. Parts of this work have been supported by the FP7 Collaborative Projects MICROME (grant agreement # 222886-2) and CEREBRAD (grant agreement # 295552), both funded by the European Commission. B.J.B. gratefully acknowledges funding from the Canadian Institutes for Health Research. CAST is freely available to academics upon request. Video production by Christos Karapiperis, background music by Kleanthis Karapiperis.

### Supplementary Materials

Supplementary Text

Methods online

Figures S1 to S2 (2)

Video S1 (1)

Tables S1 to S2 (2)

Data Files S1 to S11 (11)

References (38–63)

## References and Notes

1. Grossman, E., Medalia, O. & Zwerger, M. Functional architecture of the nuclear pore complex. Annu Rev Biophys 41, 557–84 (2012).

2. DeGrasse, J.A. et al. Evidence for a shared nuclear pore complex architecture that is conserved from the last common eukaryotic Mol Cell Proteomics 8, 2119–30 (2009).

3. Devos, D. et al. Simple fold composition and modular architecture of the nuclear pore complex. Proc Natl Acad Sci U S A 103, 2172–7 (2006).

4. Stuwe, T. et al. Architecture of the nuclear pore complex coat. Science 347, 1148–52 (2015).

5. Lee, C. & Goldberg, J. Structure of coatomer cage proteins and the relationship among COPI, COPII, and clathrin vesicle coats. Cell 142, 123–32 (2010).

6. Debler, E.W. et al. A fence-like coat for the nuclear pore membrane. Mol Cell 32, 815–26 (2008).

7. Sampathkumar, P. et al. Structure, dynamics, evolution, and function of a major scaffold component in the nuclear pore complex. Structure 21, 560–71 (2013).

8. Hoelz, A., Debler, E.W. & Blobel, G. The structure of the nuclear pore complex. Annu Rev Biochem 80, 613–43 (2011).

9. Ybe, J.A. et al. Clathrin self-assembly is mediated by a tandemly repeated superhelix. Nature 399, 371–5 (1999).

10. Brohawn, S.G., Partridge, J.R., Whittle, J.R. & Schwartz, T.U. The nuclear pore complex has entered the atomic age. Structure 17, 1156–68 (2009).

11. Whittle, J.R. & Schwartz, T.U. Structure of the Sec13-Sec16 edge element, a template for assembly of the COPII vesicle coat. J Cell Biol 190, 347–61 (2010).

12. Brohawn, S.G., Leksa, N.C., Spear, E.D., Rajashankar, K.R. & Schwartz, T.U. Structural evidence for common ancestry of the nuclear pore complex and vesicle coats. Science 322, 1369–73 (2008).

13. Wilson, K.L. & Dawson, S.C. Functional evolution of nuclear structure. J Cell Biol 195, 171–81 (2011).

14. Schwartz, T. Functional insights from studies on the structure of the nuclear pore and coat protein complexes. Cold Spring Harb Perspect Biol 5(2013).

15. Ishikawa, H. & Marshall, W.F. Ciliogenesis: building the cell’s antenna. Nat Rev Mol Cell Biol 12, 222–34 (2011).

16. van Dam, T.J. et al. Evolution of modular intraflagellar transport from a coatomer-like progenitor. Proc Natl Acad Sci U S A 110, 6943–8 (2013).

17. Jekely, G. & Arendt, D. Evolution of intraflagellar transport from coated vesicles and autogenous origin of the eukaryotic cilium. Bioessays 28, 191–8 (2006).

18. Field, M.C., Koreny, L. & Rout, M.P. Enriching the pore: splendid complexity from humble origins. Traffic 15, 141–56 (2014).

19. Katsani, K.R. et al. Functional genomics evidence unearths new moonlighting roles of outer ring coat nucleoporins. Sci Rep 4, 4655 (2014).

20. Bonasio, R. et al. Genomic comparison of the ants Camponotus floridanus and Harpegnathos saltator. Science 329, 1068–71 (2010).

21. Hsu, V.W., Lee, S.Y. & Yang, J.S. The evolving understanding of COPI vesicle formation. Nat Rev Mol Cell Biol 10, 360–4 (2009).

22. Zanetti, G., Pahuja, K.B., Studer, S., Shim, S. & Schekman, R. COPII and the regulation of protein sorting in mammals. Nat Cell Biol 14, 20–8 (2012).

23. Field, M.C. & Dacks, J.B. First and last ancestors: reconstructing evolution of the endomembrane system with ESCRTs, vesicle coat proteins, and nuclear pore complexes. Curr Opin Cell Biol 21, 4–13 (2009).

24. Field, M.C., Sali, A. & Rout, M.P. On a bender-BARs, ESCRTs, COPs, and finally getting your coat. J Cell Biol 193, 963–72 (2011).

25. Taschner, M., Bhogaraju, S. & Lorentzen, E. Architecture and function of IFT complex proteins in ciliogenesis. Differentiation 83, S12–22 (2012).

26. Kee, H.L. et al. A size-exclusion permeability barrier and nucleoporins characterize a ciliary pore complex that regulates transport into cilia. Nat Cell Biol 14, 431–7 (2012).

27. Kee, H.L. & Verhey, K.J. Molecular connections between nuclear and ciliary import processes. Cilia 2, 11 (2013).

28. Bhogaraju, S. et al. Molecular basis of tubulin transport within the cilium by IFT74 and IFT81. Science 341, 1009–12 (2013).

29. Halbritter, J. et al. Defects in the IFT-B component IFT172 cause Jeune and Mainzer-Saldino syndromes in humans. Am J Hum Genet 93, 915–25 (2013).

30. Geisert, E.E. et al. Gene expression in the mouse eye: an online resource for genetics using 103 strains of mice. Mol Vis 15, 1730–63 (2009).

31. Nery, M.F., Gonzalez, D.J. & Opazo, J.C. How to make a dolphin: molecular signature of positive selection in cetacean genome. PLoS One 8, e65491 (2013).

32. Cevik, S. et al. Active transport and diffusion barriers restrict Joubert Syndrome-associated ARL13B/ARL-13 to an Inv-like ciliary membrane subdomain. PLoS Genet 9, e1003977 (2013).

33. Cavalier-Smith, T. Predation and eukaryote cell origins: a coevolutionary perspective. Int J Biochem Cell Biol 41, 307–22 (2009).

34. Sung, C.H. & Leroux, M.R. The roles of evolutionarily conserved functional modules in cilia-related trafficking. Nat Cell Biol 15, 1387–97 (2013).

35. Mishra, R.K., Chakraborty, P., Arnaoutov, A., Fontoura, B.M. & Dasso, M. The Nup107-160 complex and gamma-TuRC regulate microtubule polymerization at kinetochores. Nat Cell Biol 12, 164–9 (2010).

36. Pascual-Garcia, P., Jeong, J. & Capelson, M. Nucleoporin Nup98 Associates with Trx/MLL and NSL Histone-Modifying Complexes and Regulates Hox Gene Expression. Cell Rep (2014).

37. Bornens, M. The centrosome in cells and organisms. Science 335, 422–6 (2012).

